# Wnts are endothelial cell-derived PKD1/PKD2-dependent autocrine/paracrine vasodilators

**DOI:** 10.64898/2026.03.17.712518

**Authors:** Ulrich C. Mbiakop, Charles E. Mackay, Alejandro Mata-Daboin, Roberto Braz Pontes, M. Dennis Leo, Jonathan H. Jaggar

## Abstract

**Background:** Wingless/Int-1 (Wnts) proteins are canonical Frizzled receptor ligands. Recent evidence indicates that some Wnts, including Wnt9b and Wnt5a, bind to polycystin 1 (PKD1), a transmembrane protein which can couple to polycystin 2 (PKD2) to form a non-selective cation channel. The functional significance of Wnts binding to PKD1 is unclear. Here, we tested the hypothesis that Wnts act through PKD1/PKD2 channels on endothelial cells (ECs) to regulate arterial contractility and blood pressure and investigated the cellular source and secretory regulation of vasoactive Wnt proteins.

**Methods:** A wide variety of approaches, including inducible EC-specific PKD1 and PKD2 knockout mice, reverse-transcription polymerase chain reaction, Western blotting, immunofluorescence, pressurized artery myography, blood pressure measurements, patch-clamp electrophysiology, *in vivo* and *in vitro* Wnt and nitric oxide assays, and Wnt secretion assays.

**Results:** Intravascular Wnt9b or Wnt5a administration stimulates an EC PKD1/PKD2-dependent dilation in pressurized resistance-size arteries. Wnt9b and Wnt5a are present in serum and plasma and intravenous infusion rapidly stimulates a blood pressure reduction which requires EC PKD1. Wnts stimulate a PKD1-dependent non-selective cation current in ECs which through Ca^2+^ signaling activates endothelial nitric oxide synthase (eNOS) and small conductance Ca^2+^-activated K^+^ channels to induce vasodilation. Wnt9b acts solely via PKD1/PKD2 channels, whereas Wnt5a stimulates signaling through PKD1/PKD2, Frizzled-7 (Fzd-7), Dishevelled and c-Jun N-terminal kinase (JNK). Intravascular flow stimulates angiotensin II type 1 (AT1) receptors, which through G_q/11_ and Porcupine activate Wnt9b and Wnt5a secretion in ECs. Wnts secreted in response to flow activate PKD1/PKD2 signaling in ECs and contribute to flow-mediated vasodilation.

**Conclusions:** Intravascular flow activates AT1 receptors, which through G_q/11_ and Porcupine stimulate Wnt9b and Wnt5a secretion in ECs. Wnt9b activates PKD1/PKD2 channels whereas Wnt5a stimulates both PKD1/PKD2 and Fzd-7 in ECs to induce vasodilation. Wnts contribute to flow-mediated autocrine/paracrine dilation and reduce blood pressure.

**Graphical abstract:** 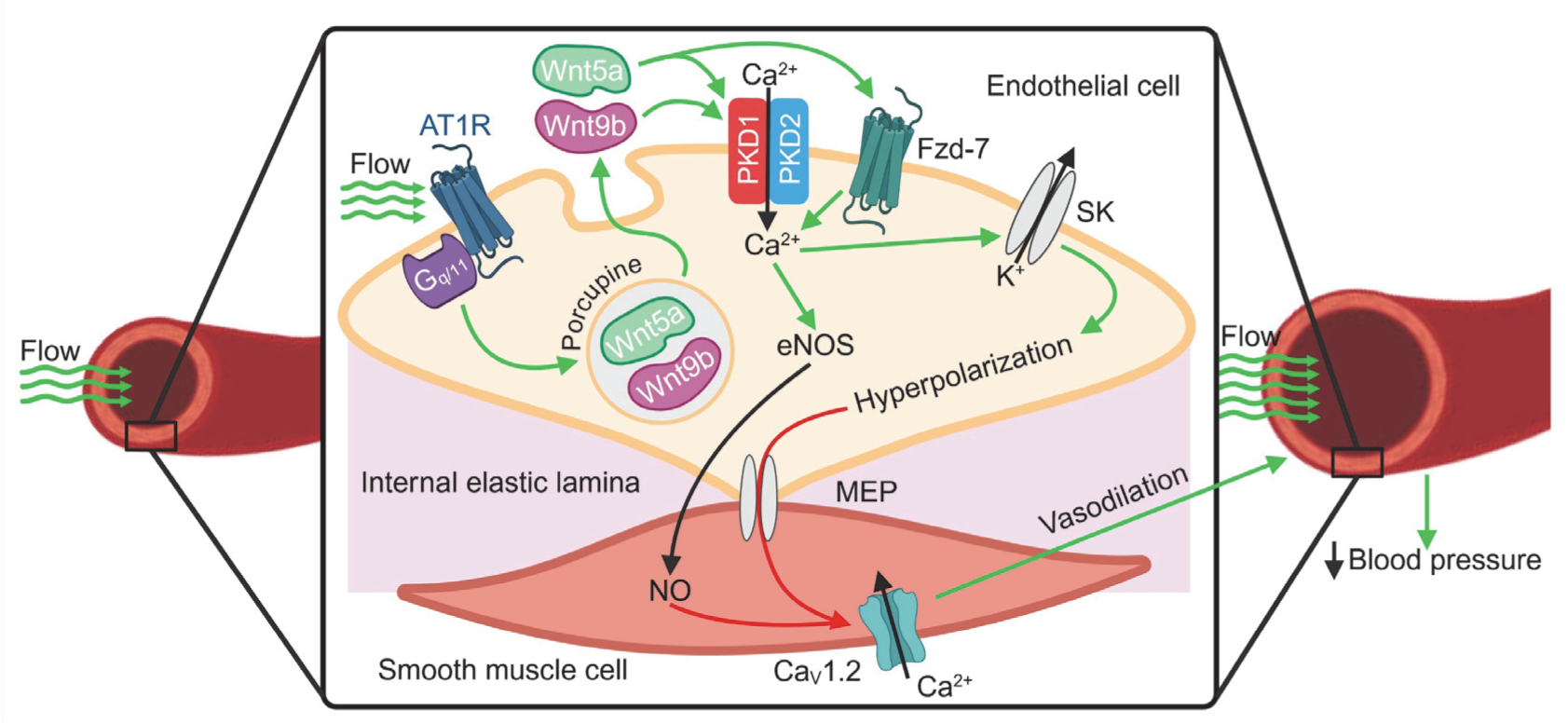

## INTRODUCTION

Wingless/Int-1 (Wnts) proteins are lipid-modified secreted glycoproteins which activate receptor-mediated signaling in cells ^1^. The Wnt family contains at least 19 members which are highly conserved across species from invertebrates to mammals ^2^. Wnts regulate cellular processes including developmental patterning, proliferation, differentiation, and polarity ^3^. Wnts classically bind to Frizzled receptors, a complex composed of Frizzled and low-density lipoprotein receptor-related protein 5 and 6 (LRP-5/6) ^2^. Wnts activate canonical β-catenin and noncanonical β-catenin-independent signaling pathways, with the latter sub-categorized into the planar polarity and Wnt/Ca^2+^ pathways ^1,4^. β-catenin-mediated Wnt signaling leads to β-catenin stabilization, leading to its nuclear translocation and the subsequent transcriptional regulation of genes which regulate proliferation, differentiation, tissue expansion, and cell fate ^5,6^. Noncanonical Wnt signaling pathways do not engage β-catenin and are considered to recruit Dishevelled ^7–10^. Noncanonical Wnt signaling can modulate processes including cell polarization, cell fate, inflammation, and cell migration ^8,11,12^. The receptors, signaling mechanisms and physiological functions of noncanonical Wnt/Ca^2+^ signaling are poorly understood. Recent evidence indicates that several Wnts can bind to polycystin 1 (PKD1, PC1), a transmembrane glycoprotein expressed by the *Pkd1* gene ^13^. In contrast to knowledge of Wnt signaling mediated by Frizzled receptors, physiological functions of Wnt signaling through PKD1 are unclear.

PKD1 contains a long extracellular N-terminus, eleven transmembrane helices and an intracellular C-terminus ^14^. PKD1 is proposed to act as a mechanosensor and ligand receptor in cultured kidney epithelial cells, although physiological stimuli which activate PKD1 are unclear ^14,15^. Physiological functions of PKD1 are also poorly understood with most knowledge related to its pathology. Noncoding and missense mutations in PKD1 lead to autosomal dominant polycystic kidney disease, which is one of the most prevalent monogenic disorders in humans^16^. PKD1 can couple to polycystin 2 (PKD2, PC2), a transient receptor potential polycystin (TRPP) channel, to form a heterotetrameric protein ^17–19^.

Several Wnts, including Wnt9b and Wnt5a, bind with low nanomolar affinity to the PKD1 N-terminus with this binding independent of Frizzled receptors ^13^. Wnts which bind to the PKD1 N-terminus activate a recombinant PKD1/PKD2 non-selective cation current in transfected CHO-K1 and *Drosophilia* S2 cells and in mouse embryonic fibroblasts^13^. Investigating Wnt-mediated PKD1 signaling in native cell types may provide important insights into the physiological functions of this signaling pathway. Endothelial cells express PKD1 and PKD2 and provide an excellent system in which to investigate Wnt-mediated signaling and function ^20–22^.

Endothelial cells regulate multiple vascular functions, including contractility. Endothelial cells regulate arterial contractility by directly controlling the membrane potential of smooth muscle cells and by producing diffusible vasodilators such as nitric oxide ^23,24^. Intravascular flow activates a plasma membrane PKD1/PKD2-dependent current in endothelial cells, leading to vasodilation and reduction in blood pressure ^21,22^. How flow stimulates PKD1/PKD2 signaling is unclear and whether Wnts regulate arterial contractility does not appear to have been explored.

Here, we tested the hypothesis that Wnts are physiological vasodilators which activate PKD1/PKD2 channels on endothelial cells, investigated the cellular source of vasoactive Wnts, and examined physiological stimuli which activate Wnt secretion to control arterial contractility. Our data indicate that intravascular flow stimulates endothelial cells to secrete Wnt9b and Wnt5a. Wnt9b activates PKD1/PKD2 channels, whereas Wnt5a activates both PKD1/PKD2 and Frizzled receptor 7 (Fzd-7) to stimulate eNOS in endothelial cells. Wnts act in an autocrine/paracrine manner and stimulate PKD1/PKD2 channels on endothelial cells to induce vasodilation and reduce blood pressure. Thus, we identify Wnts as circulating factors which induce vasodilation.

## MATERIALS AND METHODS

### Mouse models

All procedures were approved by the Animal Care and Use Committee of the University of Tennessee (#23–0444.0). *Pkd1^fl/fl^* mice with loxP sites flanking exons 2-4 of the *Pkd1* gene and *Pkd2^fl/fl^* mice with loxP sites flanking exons 11-13 of the *Pkd2* gene were obtained from the Maryland PKD Research and Translational Core Center. Cdh5(PAC)-CreERT2 mice were a kind gift from Cancer Research UK. *Pkd1^fl/fl^* mice were crossed with Cdh5(PAC)-CreERT2 mice to generate *Pkd1^fl/fl^*:Cdh5(PAC)-CreERT2 mice. *Pkd2^fl/fl^* mice were crossed with Cdh5(PAC)-CreERT2 mice to generate *Pkd2^fl/fl^*:Cdh5(PAC)-CreERT2 mice. Male *Pkd1^fl/fl^*:Cdh5(PAC)-CreERT2, *Pkd2^fl/fl^*:Cdh5(PAC)-CreERT2, *Pkd1^fl/fl^* and *Pkd2^fl/fl^* mice (12 weeks) were injected with tamoxifen (50 mg/kg, i.p.) once per day for 5 days and studied 14 days after the last injection. The genotypes of all mice were confirmed using PCR (Transnetyx, Memphis, TN) before use, as previously described^21,22^.

### Tissue preparation and endothelial cell isolation

Male *Pkd1^fl/fl^*, *Pkd1* ecKO*, Pkd2^fl/fl^* and *Pkd2* ecKO mice were euthanized with isoflurane (1.5%), followed by decapitation. Mesenteric artery branches from first- to fifth-order were removed and placed into ice-cold physiological saline solution (PSS) which contained (in mM): 112 NaCl, 6 KCl, 24 NaHCO_3_, 1.8 CaCl_2_, 1.2 MgSO_4_, 1.2 KH_2_PO_4_ and 10 glucose, gassed with 21% O_2_, 5% CO_2_ and 74% N_2_ to pH 7.4. Arteries were then cleaned of adventitial tissue. Endothelial cells were dissociated by introducing endothelial cell basal media (Endothelial cell GM MV2, Promocell) containing 2 mg/ml collagenase type 1 (Worthington Biochemical) into the lumen and incubated for 30-40 min at 37°C. Dissociated endothelial cells were maintained in ice-cold PSS and used for experimentation within 12 hours.

### Reverse transcription-polymerase chain reaction (RT-PCR)

Endothelial cells were isolated from mesenteric arteries and purified using magnetic beads (Pierce™ protein A/G magnetic beads (Thermo Fisher Scientific) coated with mouse CD31 (Abcam) primary antibody by following the manufacturer’s instructions. First-strand cDNA was synthesized from 1 to 5 ng total RNA using SuperScript IV (Invitrogen, Life Technologies). PCR was performed on first-strand cDNA using the following conditions: an initial denaturation at 94°C for 2 min, followed by 40 cycles of denaturation at 94° C for 30 s, annealing at 56°C for 30 s, and final extension at 72°C for 10 min. PCR products were separated on 2% agarose-TEA gels. Primers were used to amplify transcripts of PKD1, PKD2, myosin heavy chain 11 (Myh11), platelet-endothelial cell adhesion molecule 1 (PECAM-1) and actin (Table 1).

**Table 1.**
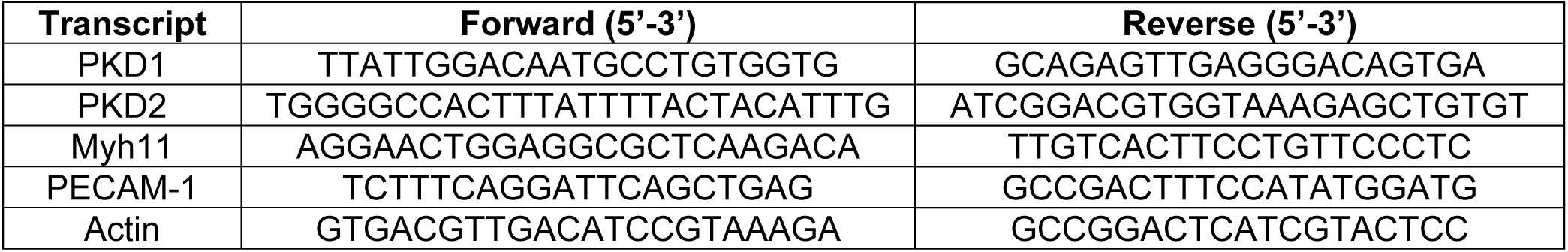
Primers used for RT-PCR.

### Western blotting

Western blotting was performed to identify proteins present in mesenteric artery (first to fifth order) or endothelial cell lysate, in the extracellular solution of arteries or endothelial cells, or in plasma or serum. For experiments measuring total endothelial nitric oxide synthase (eNOS) and phospho-eNOS (at serine 1176), the entire mesenteric artery bed from first- to fifth-order was used. A glass cannula was inserted into the first-order mesenteric artery and either Wnt9b (3μg/ml) or PSS (control) were introduced intraluminally, after which arteries were incubated for 5 min at 37°C. Mesenteric arteries or endothelial cells were transferred to ice-cold radioimmunoprecipitation assay (RIPA) buffer (Sigma-Aldrich) containing a protease and phosphatase inhibitor cocktail (1:100 dilution). The resulting lysate from mesenteric arteries or endothelial cells was centrifuged at 12,000 rpm for 10 min at 4°C, and the supernatant was assayed for protein concentration using Bradford reagent (Bio-Rad). Proteins were separated on 7.5% SDS-polyacrylamide gels and blotted onto nitrocellulose membranes. Membranes were blocked with 5% milk or 5% BSA and incubated with one of the following primary antibodies: PKD1 (Polycystic Kidney Disease Research Resource Consortium, Baltimore), PKD2 (Alomone), eNOS (Abcam), p-eNOS (Cell Signaling), Wnt9b (R&D Systems), Wnt5a (R&D Systems), AT1 receptor (Alomone) or actin (Cell Signaling) overnight at 4°C. Membranes were washed and incubated with horseradish peroxidase-conjugated secondary antibodies at room temperature for 1h. Protein bands were imaged using a ChemiDoc Touch Imaging System (Bio-Rad), quantified using ImageJ software, and normalized to actin.

### En-face arterial immunofluorescence

Arteries were cut longitudinally and fixed with 4% paraformaldehyde in PBS for 1 hr. Following a wash in PBS, arteries were permeabilized with 0.2% Triton X-100, blocked with 5% goat serum and incubated overnight with CD31 (Abcam) primary antibody and a PKD1 or PKD2 primary antibody at 4°C. Arteries were then incubated with Alexa Fluor 488 donkey anti-rat secondary antibody (1:400; Thermo Fisher Scientific), Alexa Fluor 546 rabbit anti-mouse secondary antibody (1:400; Molecular Probes) and 4’,6-diamidino-2-phenylindole, dihydrochloride (DAPI) (1:1000; Thermo Scientific) for 1 hr at room temperature. DAPI, Alexa 488 and Alexa 546 were excited at 350 nm, 488 nm and 555 nm with emission collected at 437 nm and 555 nm, respectively, using a Zeiss LSM 710 laser-scanning confocal microscope.

### Pressurized artery myography

Experiments were performed using isolated third- and fourth-order mesenteric arteries using PSS gassed with 21% O_2_/5% CO_2_/74% N_2_ (pH 7.4). Arterial segments 1-2 mm in length were cannulated at each end in a perfusion chamber (Living Systems Instrumentation) continuously perfused with PSS, and maintained at 37°C. Intravascular pressure was altered using a Servo pump model PS-200-P (Living Systems Instrumentation) and monitored using upstream and downstream pressure transducers. Following the development of stable myogenic tone, Wnt9b (R&D System), Wnt5a (R&D System), boiled (95°C, 15 min) Wnt9b, or boiled (95°C, 15 min) Wnt5a were applied intraluminally under constant flow in pressurized (80 mmHg) mesenteric arteries using a P720 peristaltic pump (Instech). Changes in arterial diameter were measured at 1 Hz using a CCD camera attached to a Nikon TS100-F microscope and the automatic edge-detection function of IonWizard software (Ionoptix). Myogenic tone was calculated as: 100 x [1-(D_active_)/(D_passive_)], where D_active_ is active arterial diameter and D_passive_ is the diameter determined in the presence of Ca^2+^-free PSS supplemented with 5 mM EGTA.

### Plasma and serum Wnt protein detection and quantification

Mice were subjected to light anesthesia with isoflurane (1.5%) and blood was collected by cardiac puncture. Blood was collected either in dry tubes to harvest serum or in EDTA-containing tubes to collect plasma. Tubes were centrifuged (3000 rpm, 15 min, 4°C) and the resulting serum or plasma was processed to reduce the abundance of highly concentrated blood proteins. Albumin and immunoglobulin were depleted in samples using Aurum Serum Protein Mini Kit, following the manufacturer’s instructions (BioRad). Samples were then passed through a protein concentrator (10K MWCO, Thermo Fisher Scientific), after which proteins were resolved using Western blotting. To quantify Wnt9b in plasma, blood samples (400-500 μl) were collected by cardiac puncture before (T0) and 5 min (T1) after i.v. Wnt9b infusion. Wnt9b proteins were measured in plasma using a mouse Wnt9b Enzyme-Linked ImmunoSorbent Assay (ELISA) kit (MyBiosource) by following the manufacturer’s instructions. A spike/recovery control was performed. 50 pg of recombinant Wnt9b was “spiked” into samples and ELISA performed. The percentage recovery of the spike was calculated as follows: 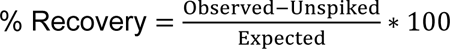; where Observed = Spiked sample value, unspiked sample value and Expected= amount spiked into the sample (calculated based on assigned concentration of spiked stock and volume spiked into the sample). To test samples for linearity, a serial dilution (1:2, 1:4, 1:8) of spiked samples were prepared and ELISA performed. Linearity was estimated as follows: 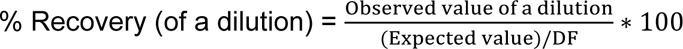; were DF = sample dilution factor A standard curve was prepared following the kit insert recommendations. A 4-parameter logistic (4PL) standard curve was generated and used to determine sample concentrations.

### Blood pressure measurement

Anesthesia was induced and maintained using 2% isoflurane. Mice were placed in a dorsal decubitus position and a catheter surgically inserted into the carotid artery. Pressure was measured using a pressure transducer coupled to a pressure monitor (PM-4, Living Systems) and a ML846 PowerLab 4/26 (AD instruments) acquisition system. Phenylephrine, prazosin, and Wnt9b were prepared in saline (0.9%) and infused into the jugular vein using a second catheter and a Hamilton syringe. Saline (0.9 %) was used as a control. Mean arterial pressure was calculated as: diastolic+⅓(systolic-diastolic).

### Nitric oxide assay

Nitric oxide was measured in cell lysate from endothelial cells and in plasma from *Pkd1^fl/fl^* and *Pkd1* ecKO mice before (T0) and 5 min (T1) after Wnt9b infusion using a nitric oxide colorimetric kit (Sigma-Aldrich), following the manufacturer’s instructions. The assay uses the enzyme nitrate reductase to convert nitrate (NO_3_) to nitrite (NO_2_). Nitrite is then detected with a microplate reader at 540 nm as a colored dye product of the Griess reaction. Total nitric oxide contributed by nitrate and nitrite in a sample is measured as nitrite after converting all nitrate to nitrite. A standard curve was obtained using recommended nitrite standard dilutions, and the nitric oxide concentration of samples was determined using the following formula: 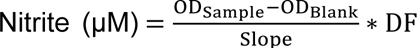. Where ODSample= optical density reading of the sample, OD_Blank_ = optical density reading of the blank, and DF = sample dilution factor.

### Cell culture and transient transfections

C57BL/6 mouse primary small intestinal microvascular endothelial cells (Cell Biologics) were studied at passages less than 5. Cells were maintained in complete endothelial cell medium containing 10% fetal bovine serum, 100 IU penicillin, and 100 μg/mL streptomycin (Cell Biologics) in a humidified incubator (5% CO_2_ and 95% air). To measure Wnt9b and Wnt5a primary antibody specificity, Wnt9b and Wnt5a were expressed in C57BL/6 mesenteric artery endothelial cells. The following vectors were transfected: pcDNA3.1 encoding full-length mouse Wnt9b (Epoch Life Science), pcDNA3.1 encoding full-length mouse Wnt5a (Epoch Life Science). Transfections were performed using Opti-MEM and Lipofectamine 2000 according to the manufacturer’s instructions (Thermo Fisher Scientific). 24 h post-transfection, culture media and cell lysate were collected for protein concentration followed by Western blotting. Angiotensin II (Ang II) type 1a and angiotensin II type 1b receptor expression were knocked down using siRNAs transfected into endothelial cells using Opti-MEM and Lipofectamine RNAiMAX according to the manufacturer’s instructions (Thermo Fisher Scientific). 24 h post-transfection, transfected cells were used for Wnt secretion assays.

For experiments measuring nitric oxide concentration, total eNOS and phospho-eNOS in C57BL/6 mouse primary small intestinal microvascular endothelial cells. Cells were cultured until confluence and were first exposed for 30 min at 37°C to either vehicle, BAPTA-AM, SRI37892 (MedChemExpress), NSC668036, (MedChemExpress) or SP600125 (MedChemExpress). Following the 30 min treatment with inhibitors, cells were exposed for an additional 5 min to either vehicle, Wnt9b or Wnt5a. Cell lysates were collected for nitric oxide assays or proteins were resolved using Western blotting.

### Patch-clamp electrophysiology

Fresh-isolated mesenteric artery endothelial cells were allowed to adhere to a glass coverslip in a recording chamber. The conventional whole cell configuration was used to measure macroscopic nonselective cation currents (I_Cat_) by applying voltage ramps between −80 mV and +80 mV from a holding potential of −40 mV. The pipette solution contained (in mM): 140 CsCl, 10 HEPES, 10 glucose, 1 EGTA, 1 MgATP, and 0.2 NaGTP (pH 7.2 adjusted with CsOH). Total Ca^2+^ was adjusted to give a final free concentration of 100 nM, whereas Mg^2+^ was adjusted to give a final free concentration of 1 mM. Free Mg^2+^ and Ca^2+^ were calculated using WebmaxC Standard (https://somapp.ucdmc.ucdavis.edu/pharmacology/bers/maxchelator/webmaxc/webmaxcS.htm). The bath solution contained 134 NaCl, 6 KCl,10 HEPES, 10 glucose, 1 MgCl_2_, and 2 CaCl_2_ (pH 7.4). Membrane currents were recorded at 5 kHZ using an Axopatch 200B amplifier, a Digidata 1332A, and Clampex 10.4 software (Molecular Devices). Currents were filtered at 1 kHz and normalized to membrane capacitance. Offline analysis was performed using Clampfit 10.4.

### Wnt secretion assay

C57BL/6 Mouse Primary Small Intestinal Microvascular Endothelial Cells (Cell Biologics) were grown to near confluence in 90mm circular dishes. On the experimental day, cells were maintained either in a static condition or under circular flow (50 rpm) for 12 h at 37°C in medium free of serum and growth factors. In some experiments, Ang II, losartan, Wnt C59, or YM 254890 were included in the culture medium. Media was then collected, centrifuged to remove any particles, and passed through a protein concentrator (Pierce™ Protein Concentrators PES, 10K MWCO, 0.5 mL, Thermo Fisher Scientific). Similar experiments were performed by exposing cannulated segments of the mesenteric arterial tree from the first- to fifth-order from *Pkd1^fl/fl^* mice to either no flow (static) or intraluminal flow (20 μl/min, 30 min) at either 4°C or 37°C, after which the PSS (extracellular solution) was collected and passed through a protein concentrator (Pierce™ Protein Concentrators PES, 10K MWCO, 0.5–100 mL). Arteries were maintained at 4°C or 37°C for 10 min prior to the introduction of flow. Western blotting was performed to detect extracellular proteins in the media or PSS and intracellular actin in the cell lysate. Extracellular proteins were then normalized to the actin of their corresponding cell lysate.

### Statistical analysis

OriginLab and GraphPad InStat software were used for statistical analyses. Values are expressed as mean ± standard error of the mean. Student t-tests were used for comparing paired and unpaired data from two populations and ANOVA with Holm-Sidak post hoc test used for multiple group comparisons. *P* values of <0.05 were considered statistically significant.

## RESULTS

### Wnt proteins stimulate an endothelial cell PKD1-dependent vasodilation and reduction in blood pressure

To investigate Wnt-mediated signaling in endothelial cells and the involvement of PKD1, we used tamoxifen-inducible endothelial cell-specific *Pkd1* knockout (*Pkd1* ecKO) mice and their genetic controls (*Pkd1^fl/fl^*). RT-PCR demonstrated that mRNA for PKD1 was present in fresh-isolated mesenteric artery endothelial cells of *Pkd1^fl/fl^* mice but was absent in endothelial cells of *Pkd1* ecKO mice (Fig. 1A). Cells used for RT-PCR expressed PECAM-1, an endothelial cell-specific marker, but did not express Myh11, a smooth muscle cell-specific marker, indicating they were pure endothelial cells (Fig. 1A). Western blotting was performed using previously validated antibodies^21,22^ and indicated that PKD1 protein in mesenteric arteries of tamoxifen-treated *Pkd1* ecKO mice was ∼57.1 % of that in tamoxifen-treated *Pkd1^fl/fl^* mice (Fig. 1B, C). This reduction in whole arterial PKD1 is expected as PKD1 is also expressed in smooth muscle cells where cre recombinase would not be activated^25^. Protein for PKD2, angiotensin II type 1 (AT1) receptors, endothelial NO synthase (eNOS) or Fzd-7, a Wnt receptor which is expressed in endothelial cells^26^, were similar in arteries of *Pkd1* ecKO and *Pkd1^fl/fl^* control mice (Fig. 1B, C). Immunofluorescence imaging demonstrated that PKD1 protein was present in endothelial cells of *en face* arteries from tamoxifen-treated Pkd1*^fl/fl^* mice but was absent in endothelial cells of tamoxifen-treated *Pkd1* ecKO mice (Fig. 1D).

**Figure 1.**
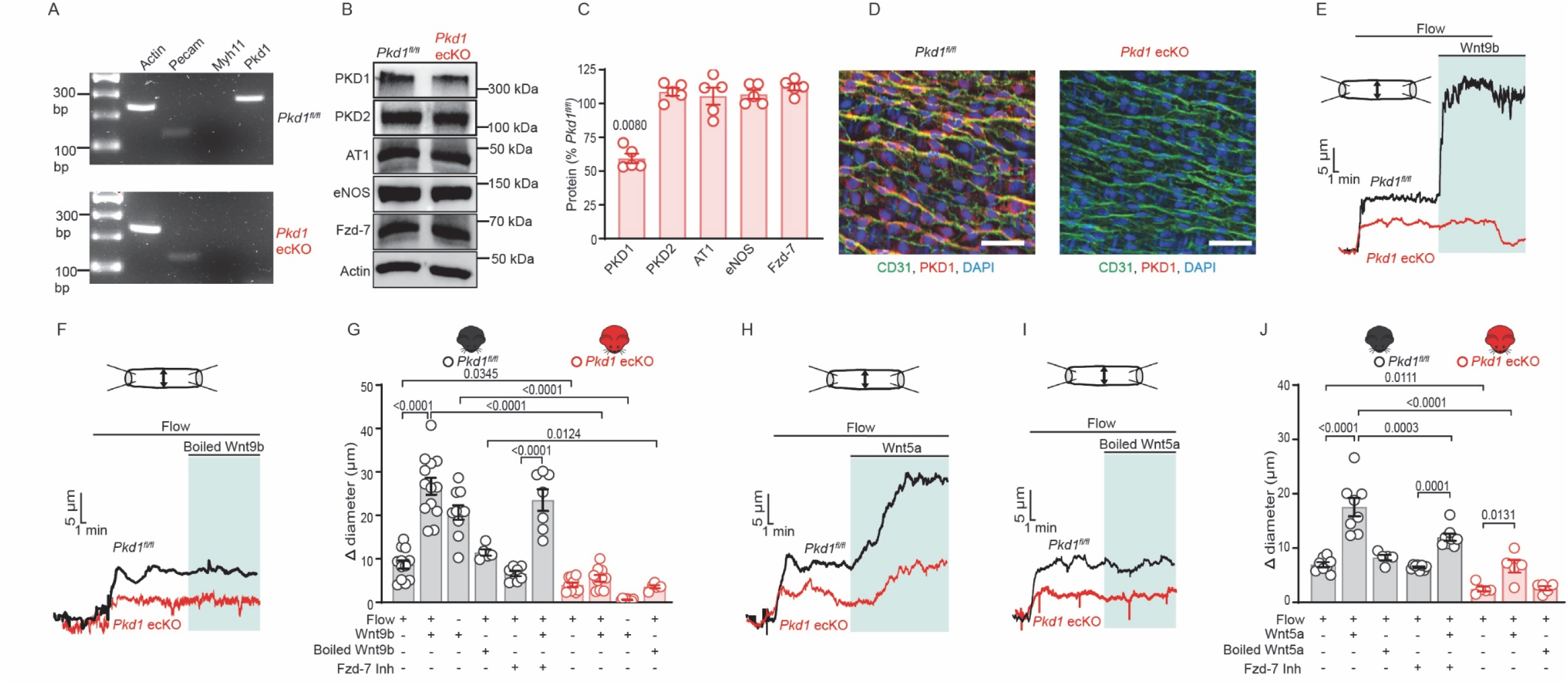
Wnt proteins stimulate an endothelial cell-PKD1-dependent vasodilation. (A) Representative RT-PCR showing that mRNA for PKD1 is present in isolated endothelial cells from tamoxifen-treated *Pkd1^fl/fl^* but absent in isolated endothelial cells from tamoxifen-treated *Pkd1^fl/fl^*:Cdh5(PAC)-CreERT2 mice. Representative of 5 independent experiments for each genotype. (B) Representative Western blots illustrating PKD1, PKD2, AT1, eNOS, Fzd-7 and actin proteins in mesenteric arteries of *Pkd1^fl/fl^* and *Pkd1* ecKO mice. (C) Mean data from experiments shown in panel B. Significance was assessed using Student t-tests, n=5 independent mesenteric arterial lysates for each genotype and each protein. (D) Immunofluorescence images (representative of three mesenteric arteries from three mice for each genotype) illustrating that PKD1 (Alexa Fluor 546) is abolished in endothelial cells of en face mesenteric arteries from *Pkd1* ecKO mice. CD31 (Alexa Fluor 488) and DAPI are also shown. Scale bars, 50 μm. (E) Diameter responses to intravascular flow (10 dyn/cm^2^) and intravascular Wnt9b (3 μg/ml) applied in continuous flow in pressurized (80 mmHg) mesenteric arteries from *Pkd1^fl/fl^* and *Pkd1* ecKO mice. (F) Diameter responses to intravascular flow (10 dyn/cm^2^) and intravascular boiled Wnt9b (3 µg/ml) applied under continuous flow (10 dyn/cm^2^) in pressurized (80 mmHg) mesenteric arteries from *Pkd1^fl/fl^* and *Pkd1* ecKO mice. (G) Mean data from experiments shown in panels E and F, and the modulation of flow and Wnt9b-mediated vasodilation by SRI37892 (Fzd-7 Inh, 5 µM). Significance was assessed using a two-way ANOVA with Holm-Sidak post hoc multiple comparisons test. n=10 arteries from 7 mice of each genotype for flow, flow+ Wnt9b and Wnt9b. n=7 arteries from 5 mice for Fzd-7 Inh+flow and Fzd-7 Inh+flow+Wnt9b. n=5 arteries from 5 mice for each genotype for flow + boiled Wnt9b. (H) Diameter responses to intravascular flow (10 dyn/cm^2^) and intravascular Wnt5a (3 µg/ml). (I) Diameter responses to intravascular flow (10 dyn/cm^2^) and intravascular boiled Wnt5a (3 µg/ml). (J) Mean data from experiments shown in panels H and I and the modulation of flow and Wnt5a-induced vasodilation by SRI37892 (Fzd-7 Inh, 5 μM). Significance was assessed using two-way ANOVA with Holm-Sidak post hoc multiple comparisons test. n=8 arteries from 6 mice from *Pkd1^fl/fl^* for flow, flow+ Wnt5a. n=5 arteries from 5 mice from *Pkd1* ecKO for flow, flow+ Wnt5a. n=5 arteries from 5 mice of each genotype for flow + boiled Wnt5a. n=8 arteries from 6 mice from *Pkd1^fl/fl^* for flow+ Fzd-7 Inh and flow+ Wnt5a+ Fzd-7 Inh.

Pressurized myogenic third-order mesenteric arteries were used to investigate the regulation of arterial contractility by Wnt9b and Wnt5a. Wnt proteins were introduced into the arterial lumen using a low rate of intravascular flow to reduce the activation of flow-mediated dilation. Low flow stimulated vasodilation in *Pkd1^fl/fl^* mouse arteries (Fig.1E-G). The subsequent luminal introduction of Wnt9b or Wnt5a under low flow produced further vasodilation in arteries of *Pkd1^fl/fl^* mice (Fig. 1E-J). Wnt-mediated dilation persisted when luminal flow was stopped in *Pkd1^fl/fl^* mouse arteries (Fig. 1E, G). Low flow stimulated a smaller dilation in *Pkd1* ecKO than *Pkd1^fl/fl^* mouse arteries (Fig.1E-J). In contrast to the vasodilation to Wnt9b which occurred in *Pkd1^fl/fl^* arteries, Wnt9b did not alter the diameter of *Pkd1* ecKO arteries under low flow or in the absence of flow (Fig. 1E-G). Dilation to Wnt5a in *Pkd1* ecKO arteries was also smaller, at ∼38.9 % of that in *Pkd1^fl/fl^* arteries when subtracting the dilation caused by low flow (Fig. 1H-J). Boiled, denatured Wnt9b and Wnt5a did not alter the diameter of *Pkd1^fl/fl^* or *Pkd1* ecKO mouse arteries (Fig. 1F, G, I, J). SRI37892, a Fzd-7 inhibitor, reduced the dilation to Wnt5a but did not alter the dilation to Wnt9b (Fig. 1G, J). SRI37892 did not alter myogenic tone or the dilation to low flow in *Pkd1^fl/fl^* mouse arteries (Fig. 1J, S1C). Depolarization (60 mM K^+^)-induced constriction, myogenic tone, vasodilation to acetylcholine (ACh), and passive diameter were all similar in *Pkd1^fl/fl^* and *Pkd1* ecKO mouse arteries (Fig. S1A-F). These data indicate that Wnt9b and Wnt5a stimulate endothelial cell PKD1-dependent vasodilation and show that Wnt5a also stimulates dilation through Fzd-7.

We investigated the hypothesis that Wnts are circulating vasodilators which are secreted by endothelial cells. To test this hypothesis, we first validated methods to detect intracellular and secreted Wnt proteins in endothelial cells. Wnt proteins undergo a variety of post-translational modifications required for secretion, including peptide cleavage, palmitoleoylation, and glycosylation ^2,3,27^. Wnt proteins also bind to Evi/Wntless (Evi/Wls), a cargo protein which is required for their surface trafficking ^28,29^. The Wnt-Evi/Wi complex can separate from Wnts prior to secretion or remain tethered to Wnts during secretion, with this process likely to be context- and cell type-dependent ^3,30,31^. Western blotting was performed to validate the Wnt9b and Wnt5a antibodies and determine the molecular weights of the secreted and intracellular Wnt9b and Wnt5a proteins in endothelial cells. In control endothelial cells, the molecular weight of intracellular Wnt9b was larger than that of extracellular Wnt9b (Fig. 2A). Transient transfection and overexpression of recombinant Wnt9b increased the amount of secreted Wnt9b protein but did not alter the abundance of intracellular Wnt9b or the molecular weights of intracellular or secreted Wnt9b (Fig. 2A, B). In contrast, the molecular weights of endothelial cell intracellular and secreted Wnt5a were similar, consistent with a previous study in mouse embryonic fibroblasts ^32,33^ (Fig. 2C). Wnt5a overexpression increased the amount of extracellular Wnt5a but did not alter the quantity of intracellular Wnt5a in endothelial cells (Fig. 2C, D). These data demonstrate that endothelial cells secrete Wnt9b and Wnt5a, identify the molecular weights of intracellular and secreted Wnt9b and Wnt5a, and validate the antibodies used to detect these proteins.

**Figure 2:**
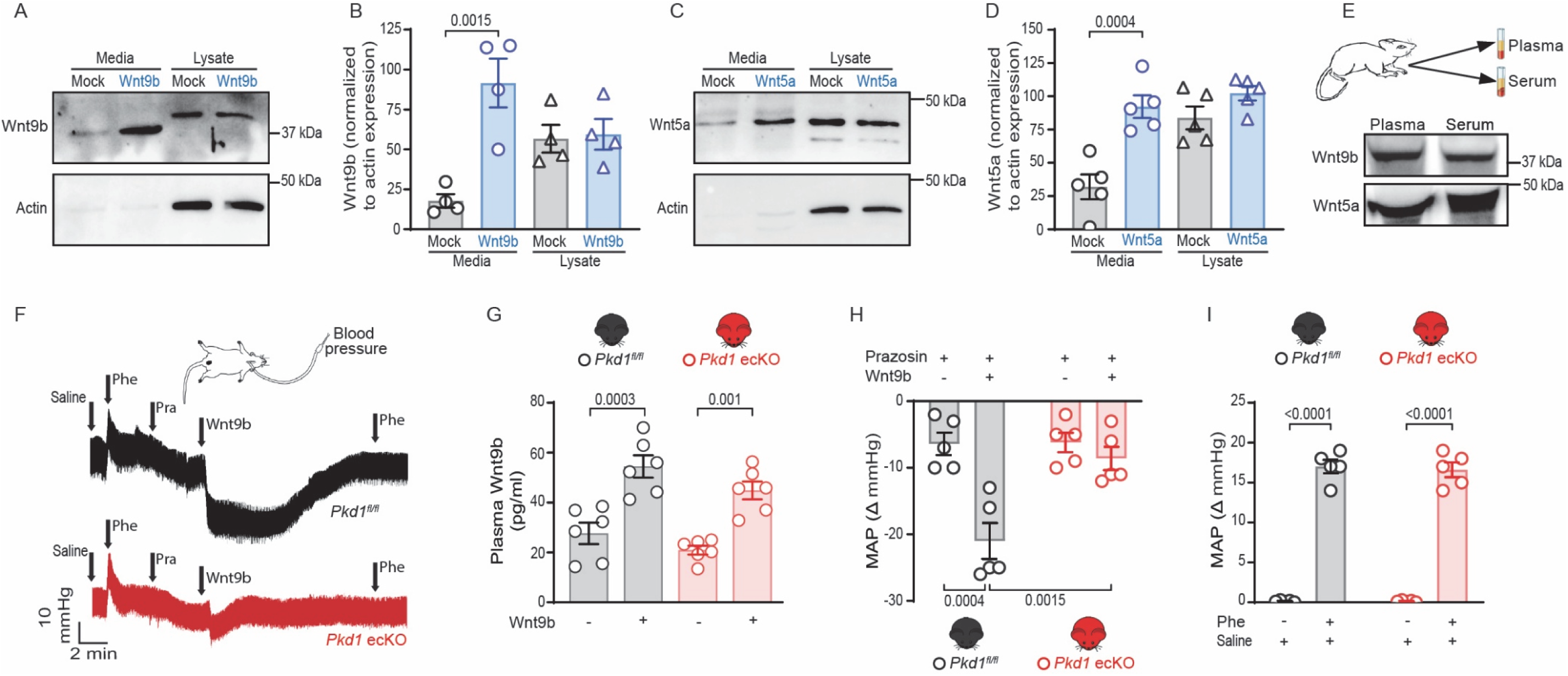
Endothelial cells secrete Wnts, which reduce blood pressure through endothelial cell PKD1. (A) Western blot of media and corresponding cell lysate from mock-transfected (mock) and Wnt9b-transfected endothelial cells illustrating secreted and intracellular Wnt9b, respectively. Representative of 4 independent experiments. (B) Mean data from experiments shown in panel A. Significance was assessed using one-way ANOVA with Holm-Sidak post hoc multiple comparisons test. (C) Western blot of extracellular solution and corresponding cell lysate illustrating secreted and intracellular Wnt5a, representative of 5 independent experiments. (D) Mean data from experiments shown in panel C. Significance was assessed using one-way ANOVA with Holm-Sidak post hoc multiple comparisons test. (E) Western blot illustrating that Wnt9b and Wnt5a are present in plasma and serum of *Pkd1^fl/fl^* mice. Representative of 3 independent experiments. (F) Representative traces illustrating the modulation of mean arterial blood pressure (MAP) by intravenous administration of saline (NaCl, 0.9 %), phenylephrine (Phe, 1 μg/kg), prazosin (Pra, 3 μg/kg) and Wnt9b (30 μg/kg) in *Pkd1^fl/fl^* and *Pkd1* ecKO mice. (G) Mean data of plasma Wnt9b at baseline and 5 min post Wnt9b infusion (30 μg/kg) in *Pkd1^fl/fl^* and *Pkd1* ecKO mice, n=6 mice for each genotype. Significance was assessed using two-way with Holm-Sidak post hoc multiple comparisons test. (H) Mean data for effects of prazosin and Wnt9b+prazosin on MAP, n=5 mice for each genotype. Significance was assessed using two-way ANOVA with Holm-Sidak post hoc multiple comparisons test. (I) Mean data for effects of saline and phenylephrine (Phe) shown in panel F, n=5 mice for each genotype. Significance was assessed using two-way ANOVA with Holm-Sidak post hoc multiple comparisons test.

Next, we tested the hypothesis that Wnts are circulating proteins which regulate blood pressure through endothelial cell PKD1. Western blotting experiments detected Wnt9b and Wnt5a in the plasma and serum of mice which were of similar molecular weights to the proteins secreted by endothelial cells (Fig. 2A-E). *In vivo* measurements were performed to measure the regulation of blood pressure by intravenous infusion of Wnt9b in anesthetized mice. Conditional endothelial cell-dependent PKD1 knockout elevates blood pressure in conscious mice by abolishing a vasodilatory signaling pathway ^22^. Isoflurane-anesthesia reduced the blood pressures of *Pkd1^fl/fl^* and *Pkd1* ecKO mice, as expected ^34^, and normalized the blood pressure difference in these lines ^22^ (Fig. S2A). Prazosin, an α_1_-adrenergic receptor blocker, was administered i.v. to depress baroreceptor sympathetic feedback. Prazosin was effective as it further reduced blood pressure and blocked blood pressure elevations to i.v. injection of phenylephrine, an α_1_-adrenergic receptor agonist (Fig. 2F, S2D, E). The plasma Wnt9b concentration in *Pkd1^fl/fl^* and *Pkd1* ecKO mice was similar at ∼27 pg/ml (Fig. 2G). Intravenous infusion of Wnt9b increased the plasma Wnt9b concentration comparably in each mouse model to ∼53 pg/ml or ∼2.0-fold (Fig. 2G). Wnt9b infusion caused a prolonged decrease in both diastolic and systolic blood pressures in *Pkd1^fl/fl^* mice, which equated to a mean arterial pressure reduction of ∼14.6 mmHg (Fig. 2F, H, S2B-C). In contrast, Wnt9b induced a transient reduction in mean arterial pressure of only ∼3.2 mmHg in *Pkd1* ecKO mice (Fig. 2F, H, S2B-C). Saline vehicle control did not alter blood pressure, phenylephrine applied prior to prazosin similarly increased blood pressure, and prazosin similarly reduced blood pressure in *Pkd1^fl/fl^* and *Pkd1* ecKO mice (Fig. 2H, I, S2B-E). These data demonstrate that Wnt9b reduces blood pressure through an endothelial cell PKD1-dependent mechanism.

### Wnt9b stimulates PKD1/PKD2 channels in endothelial cells to elicit vasodilation

PKD1 acts as atypical G protein-coupled receptor and can form a heterotetramer with PKD2 in a 1:3 ratio to form a non-selective cation channel ^17–19,35,36^. To investigate the mechanisms by which Wnts signal through PKD1 in endothelial cells, patch-clamp electrophysiology was performed. Whole-cell nonselective cation currents (I_Cat_) were recorded in fresh-isolated mesenteric artery endothelial cells of *Pkd1^fl/fl^* and *Pkd1* ecKO mice. Wnt9b increased I_Cat_ ∼3.22-fold at +80 mV and 2.68-fold at −80 mV in endothelial cells of *Pkd1^fl/fl^* mice (Fig. 3A, B). Wnt9b-activated I_Cat_ was inhibited by Gd^3+^, a non-selective cation channel blocker, in *Pkd1^fl/fl^* endothelial cells (Fig. 3A, B). In contrast, Wnt9b alone and Gd^3+^ subsequently applied in the presence of Wnt9b did not alter I_Cat_ in *Pkd1* ecKO endothelial cells (Fig. 3A, B). These data indicate that Wnt9b stimulates a PKD1-dependent I_Cat_ in endothelial cells.

**Figure 3.**
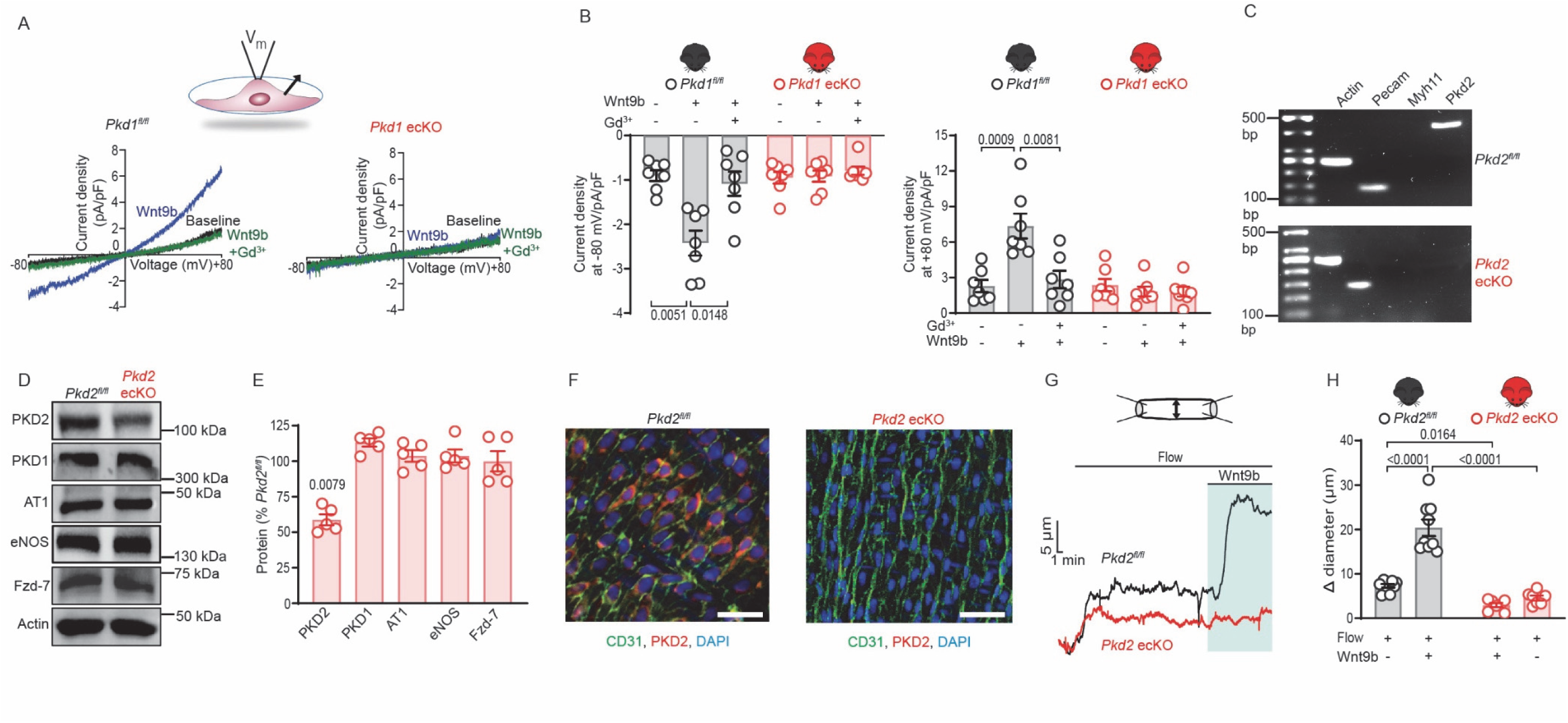
Wnt9b stimulates PKD1/PKD2 currents in endothelial cells to elicit vasodilation. (A) Representative whole-cell current recordings obtained from freshly isolated mesenteric endothelial cells of a *Pkd1^fl/fl^* and *Pkd1* ecKO mouse in control (black, Ctrl), Wnt9b (blue, 1.5 μg/ml) or Wnt9b + Gd^3+^ (green, 100 μM). (B) Mean data from experiments shown in panel A. Significance was assessed using two-way ANOVA with Holm-Sidak post hoc multiple comparisons test. n=7 cells from 5 mice from each genotype. (C) Representative gel showing that mRNA for PKD2 is present in isolated endothelial cells from tamoxifen-treated *Pkd2^fl/fl^* mice but absent in isolated endothelial cells from tamoxifen-treated *Pkd2^fl/fl^*:Cdh5(PAC)-CreERT2 mice, representative of 5 independent experiments for each genotype. (D) Representative Western blots illustrating protein levels in mesenteric arteries of *Pkd2^fl/fl^* and *Pkd2* ecKO mice. (E) Mean data from experiment shown in panel C. Significance was assessed using Student t-tests, n=5 independent mesenteric arterial lysates from each genotype and for each protein. (F) En-face immunofluorescence imaging (representative of three mesenteric arteries from three mice for each genotype) illustrating that PC-2 (Alexa Fluor 546) is present in mesenteric artery endothelial cells of *Pkd2^fl/fl^* mice, but absent in endothelial cells of *Pkd2* ecKO mice. CD31 (Alexa Fluor 488) and DAPI are also shown. Scale bars, 50 μm. (G) Diameter responses to low flow (10 dyn/cm^2^) and Wnt9b (3 µg/ml) in pressurized (80 mmHg) mesenteric arteries from *Pkd2^fl/fl^* and *Pkd2* ecKO mice. (H) Mean data for experiments shown in panel G. Significance was assessed using two-way ANOVA with Holm-Sidak post hoc multiple comparisons test. n=9 arteries from 5 mice from *Pkd2^fl/fl^* for flow, flow+ Wnt9b. n=7 arteries from 5 mice from *Pkd2* ecKO for flow, flow+ Wnt9b.

To investigate the contribution of PKD2 to Wnt-mediated vasodilation, we studied tamoxifen-inducible endothelial cell-specific PKD2 knockout (*Pkd2* ecKO) mice. RT-PCR demonstrated that mRNA for PKD2 was expressed in fresh-isolated mesenteric artery endothelial cells of *Pkd2^fl/fl^* mice but was absent in endothelial cells of *Pkd2* ecKO mice (Fig. 3C). Cells used for RT-PCR amplified transcript for PECAM-1 but did not amplify transcript for Myh11, indicating that they were pure endothelial cells (Fig. 3C). PKD2 protein in mesenteric arteries of tamoxifen-treated *Pkd2* ecKO mice was ∼63.7 % of that in tamoxifen-treated *Pkd2^fl/fl^* mice (Fig. 3D, E). The reduction in whole arterial PKD2 is expected as PKD2 is expressed in smooth muscle cells where cre recombinase would not be activated ^21^. PKD1, AT1 receptor, eNOS and Fzd-7 proteins were similar in arteries of *Pkd2^fl/fl^* and *Pkd2* ecKO mice (Fig. 3D, E). Immunofluorescence confirmed that PKD2 was present in endothelial cells of *en face Pkd2^fl/fl^* mouse arteries but absent in endothelial cells of *Pkd2* ecKO mice (Fig.3F). Intraluminal application of Wnt9b under constant low flow stimulated vasodilation in pressurized myogenic mesenteric arteries of *Pkd2^fl/fl^* mice (Fig. 3G, H). In contrast, Wnt9b did not alter the diameter of *Pkd2* ecKO mouse arteries (Fig. 3G, H). Low flow stimulated a smaller vasodilation in *Pkd2* ecKO than *Pkd2^fl/fl^* arteries, similarly to data obtained in *Pkd1* ecKO arteries (Fig. 1E-J, 3G, H). Depolarization (60 mM K^+^)-induced constriction, myogenic tone, vasodilation to ACh, and passive diameter were all similar in *Pkd2^fl/fl^* and *Pkd2* ecKO mouse arteries (Fig. S3A-F). These data indicate that Wnt9b stimulates PKD1/PKD2 channels in endothelial cells to produce vasodilation.

### Wnt9b stimulates eNOS and SK channels in endothelial cells to induce vasodilation

We investigated the intracellular signaling mechanisms by which Wnt9b stimulates vasodilation. Intraluminal application of Wnt9b under low flow stimulated a dilation which was sustained in the absence of flow in pressurized myogenic mesenteric arteries of *Pkd1^fl/fl^* mice (Fig. 4A, B). L-NNA, an eNOS inhibitor, reduced the Wnt9b-mediated dilation to ∼58.9 % of control (Fig. 4A, B). Apamin, a small conductance Ca^2+^ activated K^+^ (SK) channel inhibitor, also attenuated vasodilation to Wnt9b to ∼78.1 % of control (Fig. 4A, B). In the absence of stimuli, L-NNA and apamin do not alter the diameter of pressurized mesenteric arteries ^21^. These data indicate that Wnt9b activates eNOS and SK channels in endothelial cells to induce vasodilation.

**Figure 4.**
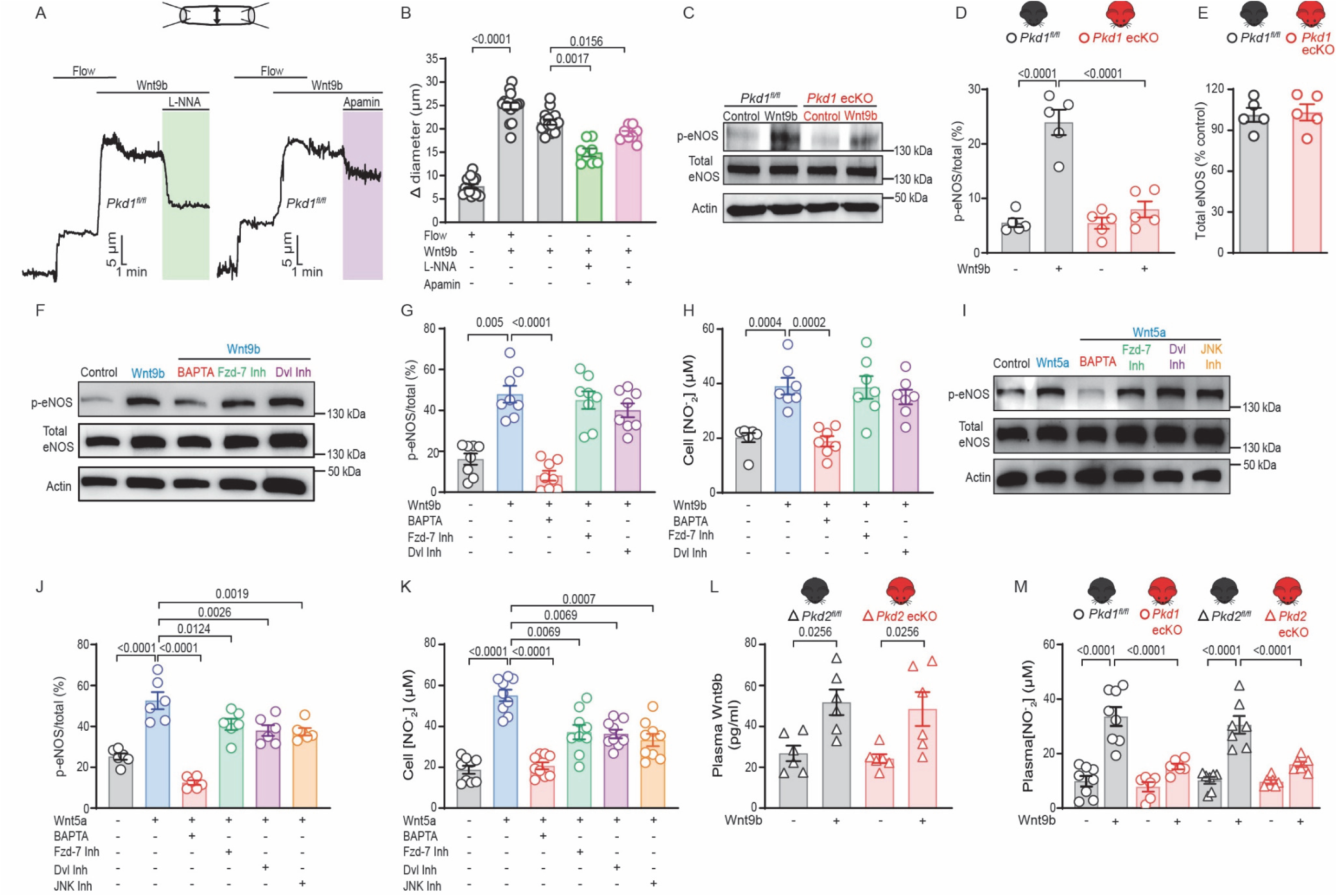
Wnt9b stimulates vasodilation through eNOS and SK channel activation. (A) Wnt9b stimulates dilation in pressurized (80 mmHg) mesenteric arteries of *Pkd1^fl/fl^* mice through eNOS and SK channel activation. Low flow (10 dyn/cm^2^) was applied and used to introduce Wnt9b (3 µg/ml) into the lumen, after which flow was stopped and L-NNA (100 µM) or apamin (300 nM) were applied abluminally. (B) Mean data for experiments shown in panel A. Significance was assessed using one-way ANOVA with Holm-Sidak post hoc multiple comparisons test. n=16 arteries from 10 *Pkd1^fl/fl^* mice for flow, flow+ Wnt9b and Wnt9b. n=8 arteries from 5 *Pkd1^fl/fl^* mice for Wnt9b+L-NNA, and Wnt9b+Apamin.(C) Western blot illustrating p-eNOS (serine1176), total eNOS, and actin in segments of first- to fifth-order mesenteric arteries of *Pkd1^fl/fl^* and *Pkd1* ecKO mice. Representative of 5 independent experiments. (D) Mean data for p-eNOS/total eNOS from experiments in panel C. Significance was assessed using one-way ANOVA with Holm-Sidak post hoc multiple comparisons test. n=5 independent mesenteric arterial lysates from each genotype (E) Mean data illustrating the effect of Wnt9b (3 µg/ml) on total eNOS from experiments in panel C. Significance was assessed using Student t-tests. n=5 independent mesenteric arterial lysates from each genotype. (F) Western blot illustrating p-eNOS (serine 1176), total eNOS, and actin in endothelial cells and modulation by Wnt9b (900 ng/ml), BAPTA-AM (BAPTA, 5 µM), SRI37892 (Fzd-7 Inh, 2.5 µM) and NSC668036 (Dvl Inh, 10 µM). Representative of 8 independent experiments. (G) Mean data from experiments shown in panel F. Significance was assessed using one-way ANOVA with Holm-Sidak post hoc multiple comparisons test. (H) Mean data illustrating nitric oxide generation from endothelial cells and modulation by Wnt9b (900 ng/ml), BAPTA-AM (BAPTA, 5 µM), SRI37892 (Fzd-7 Inh, 2.5 µM) and NSC668036 (Dvl Inh, 10 µM). Representative of 7 independent experiments. (I) Western blot illustrating p-eNOS (serine 1176), total eNOS, and actin in endothelial cells and their modulation by Wnt5a (900 ng/ml), BAPTA-AM (BAPTA, 5 µM), SRI37892 (Fzd-7 Inh, 2.5 µM), NSC668036 (Dvl Inh, 10 µM), and SP600125 (JNK Inh, 100 nM). Representative of 8 independent experiments. (J) Mean data from experiments shown in panel I. Significance was assessed using one-way ANOVA with Holm-Sidak post hoc multiple comparisons test. (K) Mean data illustrating nitric oxide generation by endothelial cells and modulation by Wnt5a (900 ng/ml), BAPTA-AM (BAPTA, 5 µM), SRI37892 (Fzd-7 Inh, 2.5 µM), NSC668036 (Dvl Inh, 10 µM), and SP600125 (JNK Inh, 100 nM). Representative of 8 independent experiments. (L) Mean data illustrating plasma Wnt9b at baseline and 5 min post Wnt9b intravascular infusion (30 μg/kg) in *Pkd2^fl/fl^* and *Pkd2* ecKO mice. n=6 mice for each genotype. Significance was assessed using two-way ANOVA with Holm-Sidak post hoc multiple comparisons test. (M) Mean data illustrating plasma nitric oxide at baseline and 5 min post Wnt9b infusion (30 μg/kg) in *Pkd1^fl/fl^*, *Pkd1* ecKO, *Pkd2^fl/fl^* and *Pkd2* ecKO mice. n=8 for *Pkd1^fl/fl^* and *Pkd1* ecKO mice. n=7 for *Pkd2^fl/fl^* and n=7 *Pkd2* ecKO mice. Significance was assessed using two-way ANOVA with Holm-Sidak post hoc multiple comparisons test.

The mesenteric arterial tree from first- to fifth-order was isolated and cannulated to measure the regulation of eNOS phosphorylation by Wnt9b. Wnt9b increased phosphorylated eNOS (p-eNOS S1176) protein ∼4.3-fold in mesenteric arteries of *Pkd1^fl/fl^* mice (Fig. 4C, D). In contrast, Wnt9b did not alter p-eNOS in *Pkd1* ecKO arteries (Fig. 4C, D). p-eNOS was similar in control in *Pkd1^fl/fl^* and *Pkd1* ecKO arteries (*Pkd1* ecKO was 98.2± 5.1 % of *Pkd1^fl/fl^*, n=5, p=0.7352) and Wnt9b did not alter total eNOS in *Pkd1^fl/fl^* or *Pkd1* ecKO arteries (Fig. 4E). Wnt9b and Wnt5a also increased p-eNOS and NO generation in endothelial cells and these effects were blocked by BAPTA-AM, a membrane-permeant Ca^2+^ chelator (Fig. 4F-K). SRI37892 or NSC668036, a Dishevelled inhibitor, did not alter the Wnt9b-induced increase in p-eNOS and NO generation in endothelial cells (Figs. 4F-H). In contrast, SRI37892, NSC668036, and SP600125, a pan- c-Jun N-terminal kinase (JNK) inhibitor, partially reduced the Wnt5a-induced increase in p-eNOS and NO generation (Fig. 4I-K). Collectively, these data indicate that Wnt9b activates PKD1/PKD2 channels, whereas Wnt5a activates signaling involving PKD1/PKD2 channels, Fzd-7, Dishevelled, and JNK in endothelial cells. Regardless of the signaling pathways, Wnt9b and Wnt5a activate Ca^2+^ signaling which stimulates eNOS phosphorylation and NO generation in endothelial cells, leading to vasodilation.

Next, we investigated whether Wnt9b-mediated PKD1/PKD2 channel signaling in endothelial cells regulates the NO concentration in plasma. Wnt9b concentrations in the plasma of *Pkd2^fl/fl^* and *Pkd2* ecKO mice were similar to those in *Pkd1^fl/fl^* and *Pkd1* ecKO mice and were comparably increased by i.v. Wnt9b infusion in all four mouse models (Fig. 2G, 4L). Intravenous Wnt9b infusion increased plasma NO in *Pkd1^fl/fl^* and *Pkd2^fl/fl^* mice but did not alter plasma NO in *Pkd1* ecKO and *Pkd2* ecKO mice (Fig. 4M). These data indicate that Wnt9b stimulates PKD1/PKD2 channels in endothelial cells, leading to eNOS activation and NO generation *in vivo*.

### Flow stimulates Wnt protein secretion in mouse mesenteric arteries and endothelial cells

Next, the cellular source of vasodilator Wnts and physiological stimuli which activate Wnt secretion from that cell type were investigated. Secreted Wnts signal through both autocrine and paracrine communication^2^. Our data indicate that Wnt9b and Wnt5a are present in the circulation (Fig. 2E, G, 4L). As such, we examined whether flow stimulates endothelial cells to secrete Wnt proteins to induce autocrine/paracrine vasodilation. For these experiments, we applied a higher shear stress (15 dyn/cm^2^) which stimulates approximately half-maximal dilation to flow ^21,22^. WIF-1 is a ∼40 kDa secreted protein which binds and inhibits Wnts^37^. Intraluminal application of WIF-1 reduced flow-mediated dilation to ∼67.9 % of control in pressurized myogenic mesenteric arteries of *Pkd1^fl/fl^* mice (Fig. 5A, B). In contrast, WIF-1 did not alter flow-mediated dilation in arteries of *Pkd1* ecKO mice (Fig. 5A, D). Flow-mediated dilation in the presence of WIF-1 in *Pkd1^fl/fl^* arteries was similar to that in *Pkd1* ecKO arteries in the absence of WIF-1 (Fig. 5A, B). These data suggest that flow stimulates endothelial cells to secrete Wnts which induce localized vasodilation through an autocrine and/or paracrine mechanism which requires PKD1 in endothelial cells.

**Figure 5.**
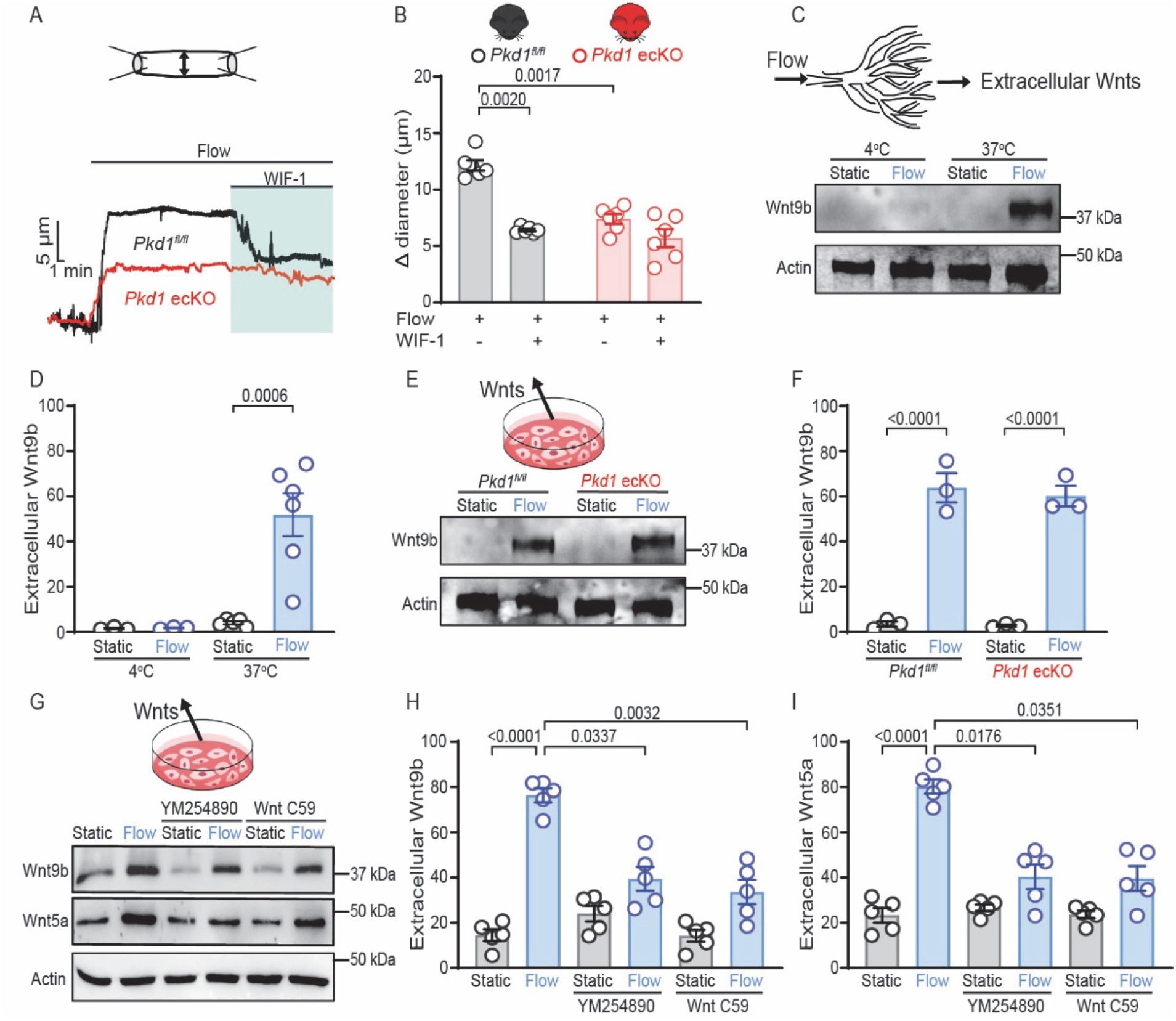
Flow stimulates endothelial cells to secrete Wnt proteins. (A) Diameter responses to intravascular flow (15 dyn/cm^2^) in the absence and presence of intraluminal WIF-1 (30 µg/ml) in pressurized (80 mmHg) mesenteric arteries from a *Pkd1^fl/fl^* and *Pkd1* ecKO mouse. (B) Mean data for experiments shown in panel A. Significance was assessed using two-way ANOVA with Holm-Sidak post hoc multiple comparisons test. n=6 arteries from 5 *Pkd1^fl/fl^* mice for flow and flow + WIF-1. n=7 arteries from 5 *Pkd1* ecKO mice for flow and flow + WIF-1. (C) Flow (20 μl/min, 30 min) stimulates Wnt9b secretion at 37°C, but not at 4°C, in mouse mesenteric arteries. Western blot of extracellular solution illustrating secreted Wnt9b and tissue lysate for corresponding actin. (D) Mean data for experiments shown in panel C with data for extracellular Wnt9b normalized to tissue actin. Significance was assessed using one-way ANOVA with Holm-Sidak post hoc multiple comparisons test. At 4°C in both static and flow conditions, n=3 independent extracellular solutions with each sample pooled from 3 mice. At 37°C, n=5 or n=6 independent extracellular solutions in static and flow conditions, respectively, with each sample pooled from 3 mice. (E) Flow (50 rpm,12 h) stimulates Wnt9b secretion similarly in primary-cultured mesenteric artery endothelial cells of *Pkd1^fl/fl^* and *Pkd1* ecKO mice. Western blot illustrating secreted Wnt9b and cell lysate for corresponding actin, representative of 3 independent experiments. (F) Mean data from experiments in panel E. Significance was assessed using one-way ANOVA with Holm-Sidak post hoc multiple comparisons test, n=3 in all groups. Wnt9b data are normalized to cellular actin. (G) Western blots of extracellular Wnt9b, extracellular Wnt5a, and cellular actin, their regulation by flow (50 rpm, 12 h) and modulation by YM 254890 (2 µM) or WntC59 (10 µM) in mesenteric artery endothelial cells. Representative of 5 independent experiments. (H, I) Mean data from experiments shown in panel G. Wnt9b and Wnt5a data are normalized to cellular actin. Significance was assessed using one-way ANOVA with Holm-Sidak post hoc multiple comparisons test. n=5 in all conditions

We tested the hypothesis that flow stimulates Wnt secretion in endothelial cells. As protein secretion is temperature-dependent, the mesenteric artery tree was cannulated and briefly (10 min) maintained at either 4°C or 37°C. This short exposure to 4°C will inhibit secretion but is not sufficient to act by blocking *de novo* protein synthesis. Arteries at 4°C or 37°C were then exposed to either intraluminal flow or no flow (static). At 37°C, flow increased extracellular Wnt9b protein 12.4±0.7-fold more than in the static condition (Fig. 5C, D). In contrast, flow did not increase extracellular Wnt9b protein at 4°C (Fig. 5C, D).

PKD1 is proposed to act as a mechanosensor ^14,15,38^. To test whether PKD1 acts as a flow sensor for Wnt secretion, mesenteric artery endothelial cells were isolated from *Pkd1^fl/fl^* and *Pkd1* ecKO mice, primary-cultured and exposed to flow or no flow at 37°C. Flow stimulated Wnt9b secretion similarly in *Pkd1^fl/fl^* and *Pkd1* ecKO endothelial cells (Fig. 5E, F). These data suggest that a PKD1-independent, flow-sensing mechanism stimulates Wnt secretion in endothelial cells.

YM 254890, a G_q/11_ inhibitor, reduced Wnt9b and Wnt5a secretion under flow to ∼43.2 % and ∼44.2 % of controls, respectively (Fig. 5G, H, I). Porcupine O-acyltransferase (PORCN) is required for Wnt palmitoylation and secretion^39–41^. WntC59, a PORCN inhibitor, also inhibited Wnt9b and Wnt5a secretion under flow to ∼56.1 % and ∼31.4 % of controls (Fig. 5G, H, I). YM254890 and WntC59 did not alter Wnt9b and Wnt5a secretion in the absence of flow (Fig. 5G, H, I). These data demonstrate that flow stimulates G_q/11_ and PORCN to induce Wnt9b and Wnt5a secretion in endothelial cells.

### Flow activates AT1 receptors to stimulate Wnt secretion in endothelial cells

Several G protein-coupled receptors are activated by shear stress, including AT1 receptors, which are expressed in endothelial cells ^42^. Shear stress and receptor ligands activate AT1 receptors by distinct mechanisms, although both mechanisms are blocked by receptor antagonists ^43–45^. Losartan, an AT1 receptor blocker, inhibited flow-mediated Wnt9b and Wnt5a secretion in primary-cultured mesenteric artery endothelial cells (Fig. 6A, B, C). In the absence of flow, losartan did not alter extracellular Wnt9b or Wnt5a (Fig. 6A, B, C). Angiotensin II, an AT1 receptor ligand, did not alter Wnt9b or Wnt5a secretion in both static and flow conditions (Fig. 6A, B, C). Transient transfection of endothelial cells with siRNAs targeting either the AT1a receptor or AT1b receptor subtype reduced AT1 receptor protein (subtype-selective AT1 receptor antibodies are not available) to ∼52.9 % and 54.4.3% of scrambled siRNA controls, respectively (Fig. S4). Flow did not alter AT1 receptor total protein in scrambled controls or in cells treated with siRNAs targeting the AT1a or AT1b receptor (Fig. S4). AT1a receptor or AT1b receptor knockdown similarly inhibited flow-activated Wnt9b and Wnt5a secretion but did not alter Wnt9b and Wnt5a secretion in a static condition (Fig. 6D, E, F). Consistent with these data, losartan reduced flow-mediated dilation in pressurized myogenic mesenteric arteries of *Pkd1^fl/fl^* mice to ∼44.8% of control but did not alter flow-mediated dilation in *Pkd1* ecKO arteries (Fig. 6G, H, I). Taken together, these data indicate that flow activates AT1 receptors which stimulate Wnt9b and Wnt5a secretion in endothelial cells, leading to a PKD1-mediated autocrine/paracrine vasodilation.

**Figure 6.**
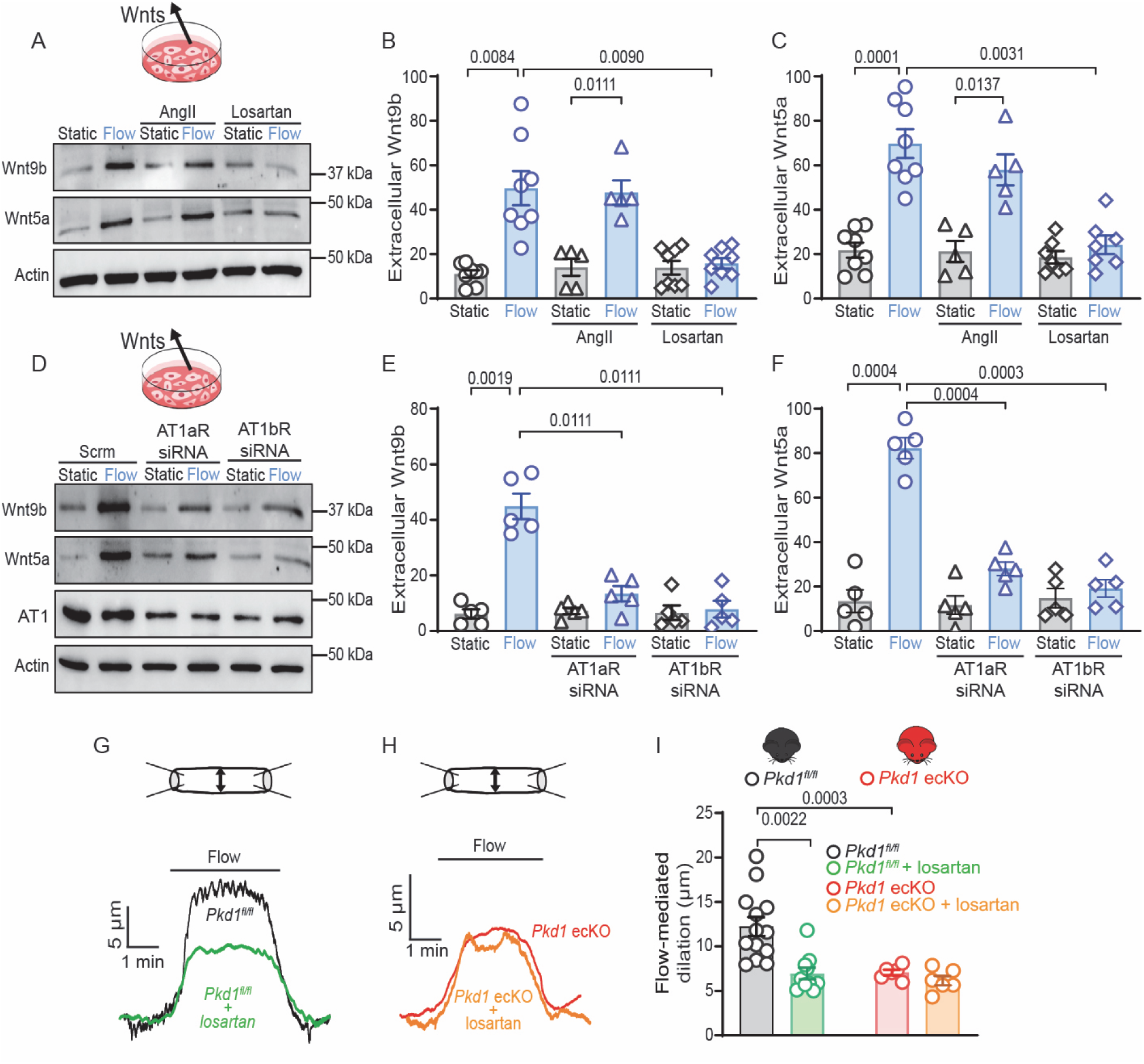
Flow activates AT1 receptors which stimulate Wnt secretion and vasodilation. (A) Western blot of extracellular solution and cellular actin illustrating that flow (50 rpm,12 h)-mediated Wnt9b and Wnt5a secretion are inhibited by losartan (1µM) in endothelial cells. In contrast, angiotensin II (0.1µM) does not stimulate Wnt secretion. Representative of 5-8 independent experiments. (B, C) Mean data for experiments shown in panel A. Wnt9b and Wnt5a data are normalized to cellular actin. Significance was assessed using one-way ANOVA with Holm-Sidak post hoc multiple comparisons test. n=8 for control, n=5 for angiotensin II, and n=8 for losartan for Wnt9b and Wnt5a in both static and flow. (D) Western blot of extracellular solution and cellular actin illustrating that flow (50 rpm, 12 h) stimulates Wnt9b and Wnt5a secretion through AT1a receptor (AT1aR) and AT1b receptor (AT1bR) activation in endothelial cells. Representative of 5 independent experiments. (E, F) Mean data for experiments shown in panel D. Significance was assessed using one-way ANOVA with Holm-Sidak post hoc multiple comparisons test. n=5 in all groups and in all conditions. (G) Flow (15 dyn/cm^2^)-mediated dilation measured in control or in losartan (1 µM) in a pressurized (80 mmHg) *Pkd1^fl/f^*l mouse mesenteric artery. (H) Diameter response to flow (15 dyn/cm^2^) measured in control or in losartan (1 µM) in a pressurized (80 mmHg) *Pkd1* ecKO mouse mesenteric artery. (I) Mean data from experiments in panels G and H. Significance was assessed using one-way ANOVA with Holm-Sidak post hoc multiple comparisons test. n=13 arteries from 10 mice from *Pkd1^fl/fl^* for flow. n=10 arteries from 7 mice from *Pkd1^fl/fl^* for flow+ losartan. n=6 arteries from 5 mice from *Pkd1* ecKO for flow, and flow+ losartan.

## DISCUSSION

Here, we investigated the regulation of arterial contractility and blood pressure by Wnts, the involvement of endothelial cell PKD1, PKD2 and Fzd-7 in mediating responses, the cellular source of vasoactive Wnts, and stimuli which activate the secretion of vasoactive Wnts. We show that Wnt9b and Wnt5a activate PKD1/PKD2 channels in endothelial cells, which through Ca^2+^ signaling stimulates eNOS and SK channels, leading to vasodilation. Wnt5a also activates Fzd-7, Dishevelled and JNK signaling to stimulate eNOS in endothelial cells. Wnt9b and Wnt5a are present in serum and plasma and Wnt9b reduces blood pressure through the activation of endothelial cell PKD1. Intravascular flow stimulates AT1 receptors, which via G_q/11_ and PORCN, stimulate endothelial cells to secrete Wnt9b and Wnt5a. Secreted Wnts contribute to flow-mediated vasodilation. Collectively, our data show that Wnt9b and Wnt5a are endothelial cell-derived circulating autocrine/paracrine factors which activate PKD1/PKD2 channels on endothelial cells to induce vasodilation and lower blood pressure.

A major finding of our study is that Wnts are a novel class of vasodilator protein. We identify endothelial cell PKD1/PKD2 channels as a principal signaling component for Wnts. To our knowledge, Wnt proteins have not previously been shown to regulate arterial contractility. Cellular processes which are known to be regulated by Wnts include embryonic development, axis patterning, cell fate specification, proliferation and migration ^3^. Wnt signaling can occur through either canonical Wnt/β-catenin-dependent or noncanonical β-catenin-independent pathways ^1,4^. Canonical Wnt signaling occurs through Frizzled receptors and a LRP-5/6 co-receptor and activates Wnt/β-catenin signaling to regulate gene expression ^2,5,6^. Noncanonical β-catenin-independent Wnt signaling is subdivided into the planar cell polarity and Wnt/Ca^2+^ pathways^1,4^. The planar cell polarity pathway uses Frizzled receptors without LRP-5/6, and can signal through Rho, a small G protein, profilin, an actin-binding protein, and rac1, a small GTPase ^46–48^. The Wnt/Ca^2+^ signaling pathway is less well defined with evidence indicating that it is stimulated by Frizzled receptors and PKD1 ^13,49,50^. Wnt9b, Wnt5a, Wnt4 and Wnt3a bind to the leucine-rich repeat and cell wall integrity and stress response component (LRR-WSC) domain of PKD1 ^13^. Wnt5a and Wnt4 also bind to the low-density lipoprotein (LDL-A) domain of PKD1 ^13^. Wnt binding to PKD1 activates a PKD1/PKD2 current in transfected CHO-K1 and *Drosophilia* S2 cells and in mouse embryonic fibroblasts ^13^. We show that Wnt9b activates a PKD1-dependent non-selective cation current in endothelial cells, stimulates eNOS via a PKD1/PKD2- and Ca^2+^-dependent mechanism, induces a PKD1/PKD2-dependent vasodilation which requires eNOS and SK channels, and stimulates an endothelial cell PKD1-dependent reduction in blood pressure. Wnt5a also stimulates eNOS via a Ca^2+^-dependent mechanism and induces an endothelial cell PKD1-dependent vasodilation. PKD1/PKD2 is Ca^2+^-permeant, suggesting that Wnt9b and Wnt5a stimulate Ca^2+^ influx through these channels to activate eNOS and SK in endothelial cells and induce vasodilation ^18,19^. Thus, Wnt9b and Wnt5a activate the Wnt/Ca^2+^ signaling pathway in endothelial cells. We show that Wnt5a also activates eNOS via Fzd-7-, Dishevelled-, and JNK-dependent signaling in endothelial cells and stimulates vasodilation through Fzd-7. It remains to be determined whether the Fzd-7 signaling pathway for Wnt5a is through canonical and/or non-canonical Wnt signaling. Opposing effects of Wnt5a on eNOS activity have previously been described in other cell types. Wnt5a inhibited insulin- or A23187 (a Ca^2+^ ionophore) induced eNOS activation through a JNK-mediated mechanism in cultured human aortic and adipose tissue endothelial cells ^51,52^. In contrast, Wnt5a stimulated NO generation in hippocampal neurons ^53^. In summary, our data indicate that Wnt9b activates PKD1/PKD2 channels, whereas Wnt5a activates both PKD1/PKD2 and Frizzled receptor 7 (Fzd-7) in endothelial cells.

We demonstrate that intravascular infusion of Wnt9b produces a sustained reduction in blood pressure which requires PKD1 in endothelial cells, similarly to the Wnt9b-mediated vasodilation in pressurized arteries. The Wnt9b concentration in blood does not appear to have previously been measured, although Wnt5a and Wnt3a are between 0.32 and 15 ng/ml in humans ^54–56^. We found plasma Wnt9b to be ∼27 pg/ml in mice. Elevating plasma Wnt9b ∼2.0-fold caused a PKD1/PKD2-dependent increase in plasma NO and a reduction in blood pressure. These data indicate that Wnt9b stimulates PKD1/PKD2 channels in endothelial cells, leading to eNOS activation and NO generation *in vivo*. We previously used radiotelemetry to show that conditional endothelial cell PKD1 or PKD2 knockout elevates blood pressure in freely-moving, conscious mice ^21,22^. The higher blood pressure in *Pkd1* ecKO and *Pkd2* ecKO mice is consistent with loss of the endothelial cell Wnt-PKD1/PKD2 pathway of vasodilation that we describe here. Here, we used anesthetized mice to measure plasma Wnt9b, plasma NO and blood pressure, and to administer Wnt9b i.v.. Isoflurane anesthesia lowers blood pressure ^34^. We found that the isoflurane-induced reduction in blood pressure also normalizes the blood pressure difference between *Pkd1^fl/fl^* and *Pkd1* ecKO mice. This blood pressure normalization is beneficial as we did not need to account for different baseline blood pressures in *Pkd1^fl/fl^* and *Pkd1* ecKO mice prior to measuring responses to i.v. Wnt9b infusion. We also found that baseline plasma NO is similar in anesthetized *Pkd1^fl/fl^*, *Pkd1* ecKO, *Pkd2^fl/fl^*, and *Pkd2* ecKO mice. There are several explanations for this finding, including that baseline concentrations of Wnt proteins release nitric oxide locally and preferentially to smooth muscle cells rather than into the circulation and that isofluorane anesthesia and the associated reduction in blood pressure and flow masks differences in plasma NO in the mouse lines. Wnt9b infusion caused a transient, small reduction in blood pressure in *Pkd1* ecKO mice, suggesting it activated PKD1-dependent or -independent signaling in other cell types. Intraventricular infusion of Wnt3a into the nucleus tractus solitarius lowered blood pressure, decreased heart rate, and increased NO release, providing one potential alternative mechanism of action for Wnt9b ^57^. Future studies should investigate additional mechanisms by which circulating Wnts induce hypotension.

Our data demonstrate that a signaling mechanism mediated by AT1 receptors, G_q/11_ and PORCN underlies flow-mediated Wnt9b and Wnt5a secretion in endothelial cells. In contrast, PKD1/PKD2 channels do not regulate Wnt9b secretion. Previous studies investigated the mechanisms by which membrane stretch activates AT1 receptors and other G protein-coupled receptors. Stretch activates AT1 receptors by causing an anticlockwise rotation and movement of the seventh transmembrane domain (TM7) into the ligand-binding pocket ^58^. Candesartan, an inverse agonist, binds to G257 and T287 in AT1 receptors, stabilizes an inactive confirmation, and prevents stretch-activated helical movement of TM7 ^58^. TM6 may also be mechanosensitive in G protein-coupled receptors ^59^ and TM8 is an essential structural component of mechanosensitive G protein-coupled receptors which leads to intracellular signaling ^60^. AT1 receptors are G_q/11_-coupled, consistent with our data demonstrating that this G protein contributes to flow-mediated Wnt secretion. We demonstrate that losartan, which is structurally similar to candesartan, inhibits flow-mediated Wnt9b and Wnt5a secretion in endothelial cells. We also show that losartan inhibits flow-mediated dilation, consistent with flow activating AT1 receptors to stimulate Wnt secretion that leads to PKD1/PKD2-mediated dilation. Mechanical stretch of AT1 receptors also activates β-arrestin signaling in the absence of ligand binding or G protein activation ^44,61^.The involvement of β-arrestin in AT1 receptor-mediated Wnt secretion in endothelial cells remains to be established. AT1 receptors can exist as both monomers, homodimers and heterodimers ^62^.

Rodents express both AT1a and AT1b receptors, whereas humans express one AT1 receptor type. We found that the knockdown of either AT1a or AT1b receptors similarly attenuated Wnt9b and Wnt5a secretion, with each causing a reduction of more than 50%. These data suggest AT1a and AT1b receptors form a flow-sensitive heterodimer in endothelial cells. In contrast, angiotensin II did not stimulate Wnt secretion, consistent with previous evidence that membrane stretch and ligands activate AT1 receptors through distinct mechanisms ^63,64^. WIF-1 reduced flow-mediated dilation in *Pkd1^fl/fl^* arteries to that in *Pkd1* ecKO arteries, supporting other evidence here that flow stimulates the secretion of PKD1-dependent autocrine/paracrine vasodilatory Wnts. Endothelial cell PKD1 knockout reduced dilation to low flow, Wnt9b and Wnt5a, whereas SRI37892 reduced dilation to Wnt5a, but did not alter dilation to low flow. Thus, the low flow we used to introduce Wnt proteins into the lumen stimulates the secretion of Wnts which activate PKD1 but not Fzd-7. Endothelial cells may release a wide variety of Wnt proteins to produce dilation. We show that WIF-1 attenuates the dilation to a higher flow rate which produces half-maximal dilation ^21,22^. These data suggest that physiological concentrations of Wnt proteins released by endothelial cells in response to flow activate PKD1/PKD2-mediated dilation. Future studies should determine the functional contribution of different Wnts, PKD1/PKD2 channels and Frizzled receptor subtypes to flow-mediated dilation at different levels of shear stress.

Previous studies demonstrated that PKD1 can contribute to mechanical signaling, leading to the proposal that PKD1 may be a mechanosensitive protein ^15,65–67^. Our results suggest that shear stress does not directly activate PKD1/PKD2 channels to elicit dilation. It is possible that higher levels of shear stress than we studied activate PKD1/PKD2 channels independently of Wnt-mediated signaling in endothelial cells. Previous studies demonstrated that the activation of Piezo1, a mechanosensitive channel expressed in endothelial cells, leads to either vasodilation, opposition of vasodilation, or vasoconstriction ^68–70^. Whether Piezo1 contributes to or modifies Wnt-PKD1/PKD2 signaling, and at what level, remains to be determined. We studied the functional significance of Wnt and PKD1/PKD2 signaling under laminar flow, which is considered to vasoprotective, in part by reducing the production of reactive oxygen species (ROS) and by increasing bioavailable NO ^71–73^. Future studies should investigate the regulation of Wnt-PKD1/PKD2 channel signaling during oscillatory or disturbed flow, which stimulates ROS production, reduces bioavailable NO, and is associated with endothelial cell dysfunction ^71–73^.

The human circulation contains approximately one trillion (10^12^) endothelial cells which are constantly exposed to flow, suggesting that each vascular endothelial cell would need to secrete only a small amount of Wnts to contribute to the circulating concentration. It remains to be determined whether endothelial cells in some organs contribute more to the circulating concentration of vasoregulatory Wnts than others. For example, liver endothelial cells are a prominent source of Wnt9b expression and may contribute more than endothelial cells in other organs to circulating Wnts ^74^. Also unclear is the expression level of PKD1/PKD2 channels in endothelial cells of different organs and the effectiveness of Wnts to induce vasodilation and reduce blood pressure in different vascular beds. Taken together, our observations suggest that endothelial cell-derived Wnt proteins stimulate vasodilation through paracrine signaling. Wnt proteins may also act through autocrine signaling as flow stimulates cation currents in isolated endothelial cells of *Pkd1^fl/fl^* and *Pkd2^fl/fl^* mice which are smaller in endothelial cells of *Pkd1* ecKO and *Pkd2* ecKO mice ^21,22^. Thus, endothelial cell-derived Wnts appear to stimulate vasodilation through both autocrine and paracrine signaling.

A logical extension of our study would be to investigate whether sexual dimorphism exists in Wnt signaling mediated by PKD1/PKD2 channels, Frizzled proteins, Dishevelled, and JNK in endothelial cells and the regulation of arterial contractility and blood pressure by these multi-modal pathways. Sexual dimorphism may also exist in the mechanisms by which intravascular flow activates Wnt secretion in endothelial cells, which types of Wnt proteins are secreted, and the circulating concentrations of different Wnts. Wnt secretion and signaling may also become dysfunctional during diseases. For example, Wnt5a is higher in the serum of patients with sepsis, a disease which is associated with a dangerous drop in blood pressure due to vasodilation ^55^. It is possible that the vasodilation during sepsis occurs in part due to Wnt-mediated PKD1/PKD2 channel activation in endothelial cells. Future studies should investigate these many possibilities.

In summary, we show that intravascular flow stimulates endothelial cells to secrete Wnt9b and Wnt5a. Wnt9b activates PKD1/PKD2 channels whereas Wnt5a activates both PKD1/PKD2 and Fzd-7 to stimulate eNOS in endothelial cells. Wnts act in an autocrine/paracrine manner to stimulate vasodilation and reduce blood pressure.

## Acknowledgments

We thank Tessa Garrud for helpful comments on the manuscript. The graphical abstract was created using Biorender.

